# The extended Split-ORF pipeline: Prediction and evaluation of Split-ORFs using Ribo-seq data

**DOI:** 10.64898/2026.05.22.727176

**Authors:** Christina Kalk, Justin Murtagh, Vladimir Despic, Michaela Müller-McNicoll, Marcel H. Schulz

## Abstract

Split Open Reading frames (Split-ORFs) occur in transcripts containing at least two open reading frames, each encoding a part of the same full-length protein. These multiple open reading frames arise from alternatively spliced transcript isoforms. Understanding which genes make Split-ORFs, and in which cell types and under which conditions, would generate new insights into gene regulation. We previously published the Split-ORF pipeline, a computational tool that predicts candidate Split-ORFs from transcript sequences. Here, we present a new and improved version of the Split-ORF pipeline adding modules to analyze Ribo-seq data, calculate regions unique to the Split-ORF candidates, quantitatively assess Ribo-seq coverage in these regions, and perform candidate prioritization. Using this pipeline, we predicted more than 14,000 candidate Split-ORF transcripts from alternatively spliced human transcripts containing premature termination codons or retained introns. Hundreds of candidate Split-ORFs show significant Ribo-seq coverage across diverse cell types and diseases in at least one of the Split-ORFs, and 120 transcripts in both Split-ORFs. The candidate Split-ORF genes with significant Ribo-seq coverage are enriched for RNA-binding and RNA-processing functions and the majority of them encode RNA-binding proteins.

## Introduction

Alternative splicing enables a single gene to produce multiple transcript and protein isoforms. In addition to increasing transcriptome and proteome diversity, alternative splicing also contributes to the tight regulation of protein and mRNA abundance. For example, when protein levels are elevated splicing events that introduce a premature termination codon into the open reading frame (ORF) are favored. The resulting premature termination codon-containing transcripts encode truncated proteins and are typically recognized and degraded by nonsense-mediated decay (NMD) (1; 2; 3). Dysregulation of alternative splicing and NMD has been implicated in various diseases, including cardiomyopathies and gastric, myeloid, and breast cancers (4; 5; 6; 7). In these contexts, translation of the aberrantly spliced transcripts can generate neoantigens, which represent promising therapeutic targets for immunotherapies in different cancer types (8; 9; 10).

Various RNA-binding proteins (RBPs), including the serine- and arginine-rich splicing factors and heterogeneous nuclear ribonucleoproteins, use alternative splicing to control their own protein abundance (11). Serine- and arginine-rich splicing factors promote the inclusion of poison cassette exons or the retention of introns, thereby introducing premature termination codons and rendering their transcripts sensitive to degradation by NMD (12). However, we recently showed that overexpression of SRSF7, which is frequently observed in cancers, such as colon and lung cancer (13; 14; 15), inhibits NMD in cis, allowing poison cassette exon-containing isoforms to accumulate. Translation of these transcripts produces a truncated SRSF7 variant that is RNA-binding competent but defective in splicing (16). Interestingly, we also identified a second start codon within the poison cassette exon, downstream of the premature termination codon, whose translation gave rise to a second truncated protein, which is in-frame with the original SRSF7 protein sequence. Because these two ORFs encode parts of the full length SRSF7 protein - effectively “splitting” it in two halves - we termed them “Split-ORFs”, see Fig. 1A-B. We showed that usage of the second start codon is required for NMD escape, and proposed that translation of the second Split-ORF removes exon-junction complexes from the transcript, thereby stabilizing it. However, the mechanism of Split-ORF translation remains elusive.

**Figure 1.**
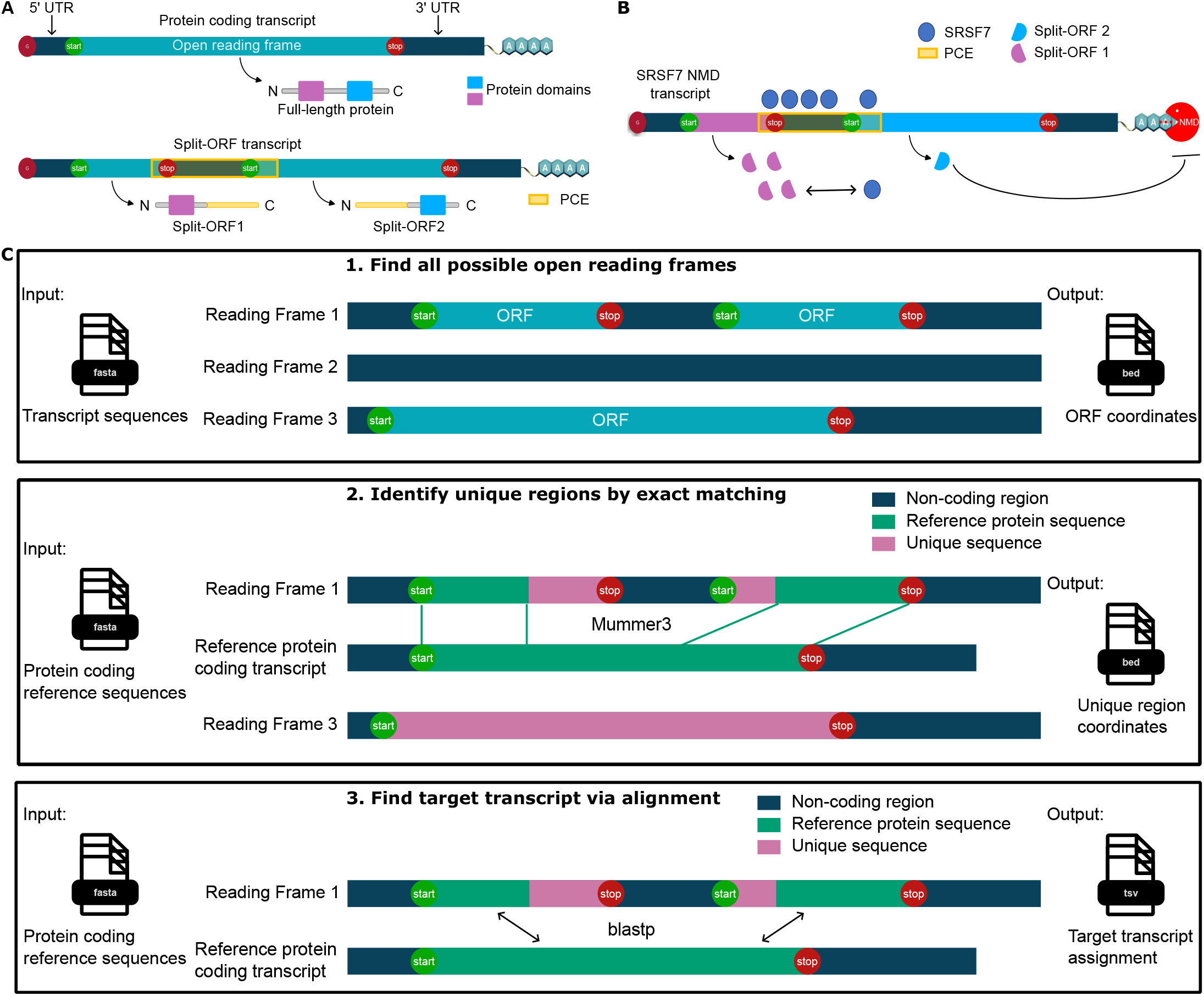
The Split-ORF pipeline predicts Split-ORFs based on transcripts’ sequences. PCE: poison cassette exon. **A**: Schematic of a Split-ORF transcript (lower transcript) with its protein-coding target transcript (upper transcript). **B**: SRSF7 forms Split-ORFs upon overexpression by binding to its own transcript. The first Split-ORF protein is a dominant-negative isoform of the full-length protein. The second Split-ORF protein renders the transcript NMD resistant. **C**: The three main steps of the Split-ORF pipeline. In the first step, all possible ORFs within the transcript’s sequence are found. In the second step, unique regions are identified by exact matching to a protein-coding reference. In the last step, the target transcript is identified via alignment to the protein-coding reference.

These findings raised the possibility that Split-ORFs and autoregulatory feedback mechanisms are more widespread than previously recognized. To test this, we developed the first version Split-ORF pipeline which predicts candidate Split-ORFs and Split-ORF transcripts from annotated NMD-sensitive transcript sequences (16). These candidate Split-ORFs require further assessment to identify high-confidence candidates for functional analyses. In our first attempt, coordinates of poison cassette exons introducing premature termination codons were retrieved manually and intersected with Ribo-seq reads. If two or more reads mapped within the poison cassette exon, the Split-ORF transcript was considered as an interesting candidate. With this analysis, we identified 19 human RBPs predicted to form Split-ORFs that fulfilled this criterium. These findings, supported by further experimentation, led to the hypothesis that Split-ORF formation might be a shared autoregulatory mechanism among a subset of RBPs.

The approach of the first version of the Split-ORF pipeline had several limitations: First, the manual retrieval of the poison cassette exon coordinates made the analysis labor-intensive and difficult to apply consistently across different datasets. Second, only Split-ORFs arising from poison cassette exons were considered. Other sources of structural variation, such as alternative splice donor and acceptor sites or the retention of introns, can also generate Split-ORFs but were excluded by design. Moreover, because candidate regions were retrieved by manual coordinate look-up against the reference annotation, only Split-ORFs formed by annotated poison cassette exons were accessible, and unannotated events were missed entirely. Third, requiring a minimum of two Ribo-seq reads mapping within the poison cassette exon is prone to false positive identification caused by mapping artifacts and other sources of noise. Furthermore, observing Ribo-seq coverage anywhere within the poison cassette exon makes it difficult to draw inference about which Split-ORF could be translated, and it does not test whether the Ribo-seq coverage is present within the Split-ORF, so the possibility remains that the reads map at positions outside of the Split-ORF coordinates.

To address these shortcoming, we added four new features to the pipeline: we determine regions that are unique to the predicted candidate Split-ORFs, we automated the usage of Ribo-seq data to evaluate the predicted candidate Split-ORFs, we perform statistical testing of Ribo-seq coverage in the unique regions against untranslated background, providing p- and q-values for the tested regions and we classify the candidate Split-ORFs into different confidence sets based on their Ribo-seq coverage status. We also produce several reports for easy use and documentation. This new and improved pipeline is referred to as the Split-ORF pipeline v2, or the extended Split-ORF pipeline, interchangeably.

We applied the Split-ORF pipeline v2 to thousands of alternatively spliced human transcripts and Ribo-seq datasets from diverse cell types and conditions, and identified hundreds of candidate Split-ORFs with significant ribosome coverage in one of the candidate Split-ORFs, as well as in both candidate Split-ORFs in a total of 120 candidate Split-ORF transcripts. The majority of interesting Split-ORF candidates are predicted to encode truncated RBPs, a subset of which showed increased ribosome coverage upon NMD inhibition. Split-ORFs were also recovered in fibrocystic disease, breast cancer and glioblastoma cell lines, consistent with NMD, Split-ORF formation and translation being regulated in a cell type- and disease-specific manner (17; 18).

## Materials and Methods

### Definition of Split-ORFs and Split-ORF transcripts

We define a Split-ORF transcript as a transcript which contains two or more ORFs, each encoding a part of the original full-length protein, as illustrated in Fig. 1A. The Split-ORFs are the respective ORFs that are translated into partial proteins. We further define unique regions as sequences which appear exclusively in the Split-ORFs. Consequently, they are not present in any other protein-coding transcript.

### The extended Split-ORF pipeline

The Split-ORF pipeline v2 predicts whether a given transcript can form a Split-ORF based on its nucleotide sequence; this we define as a candidate Split-ORF transcript. Furthermore, the pipeline calculates unique regions as DNA and protein sequences. The unique DNA sequences can be used to test for the significance of the coverage from Ribo-seq data relative to a background of untranslated regions. The pipeline is available as the `split-orf` conda package and contains two modules, called `split-orf-prediction` and `ribo-cov`. Input to the `split-orf-prediction` module are the sequences of the transcripts of interest as FASTA files, for example alternatively spliced transcripts annotated as NMD or retained intron (RI) from Ensembl, as well as sequences of reference protein-coding transcripts as FASTA files including coordinates of their coding sequences and protein domain annotations based on the protein family (PFAM) database (19) as BED and TSV files, respectively. We also used snoRNA, snRNA, scaRNA and miRNA as well as mt-rRNA and -tRNA, rRNA and rRNA-pseudogene annotations for filtering; these coordinates were downloaded from Ensmebl Biomart (v110) (20).

The file formats for in- and outputs to the Split-ORF pipeline are explained in detail in the Supplementary Information (Pipeline implementation). An overview of the steps of the Split-ORF pipeline v2 in comparison to the previous pipeline version is given in Table 1.

**Table 1.** Overview of changes in the steps of the second version of the Split-ORF pipeline compared to the steps of the first version.

| Module | Step | Split-ORF pipeline v1 | Split-ORF pipeline v2 | Status |
| --- | --- | --- | --- | --- |
| split-orf-prediction | ORF prediction | python script | python script | unchanged |
| split-orf-prediction | unique region calculation | not present | Mummer3 exact matching | new |
| split-orf-prediction | target transcript identification | alignment of ORF | alignment of non-unique regions | modified |
| ribo-cov | background calculation | not present | Ribo-seq coverage distribution in 3' UTRs | new |
| ribo-cov | p- and q-values for unique region coverage | not present | quantile of background distribution | new |
| ribo-cov | ribo-cov categorization based on ORF position | not present | unique region present in first, second or both ORFs | new |

### Steps of the split-orf-prediction module

#### Step 1: Find all candidate ORFs larger than 50 amino acids

First, all candidate ORFs of the transcripts of interest are identified from their nucleotide sequences (Fig. 1C, step 1). To achieve this, all canonical start (AUG) and stop codons (TAG, TGA, TAA) of the three reading frames of the transcript are detected. For a reading frame, the most upstream start codon is identified and an ORF defined which ends at the first in-frame stop codon. All in-frame start codons which are located between the chosen start and stop codon are discarded. The most 5’ in-frame start codon positioned downstream of the previous stop codon is selected and the procedure is repeated until no start or stop codons are left. Then all detected ORFs are filtered for a minimal length of 50 amino acids (AA). This step is implemented in a custom python script.

#### Step 2: Identify unique regions in candidate Split-ORFs by exact matching

For candidate ORFs identified in step 1, unique regions are determined. To this end, exact matches between the sequences of the ORFs and all protein-coding reference sequences are retrieved using Mummer3 (21) both on the RNA and protein sequence level. The minimum length of the exact matches is 20 bp or 8 AA for the DNA and protein sequences, respectively. Regions of the ORFs that do not match sequences of the protein-coding reference transcripts are considered unique regions (Fig. 1C, second step).

The identified unique regions are filtered for a length greater than or equal to the Mummer3 minimum match length parameter. Furthermore, unique regions located at the start of the most 5’ Split-ORF with respect to the transcript or at the end of the most 3’ Split-ORF are filtered out. These unique regions do not necessarily constitute evidence for Split-ORF formation, as they could be explained by alternative start and stop codons. Because the Mummer3 exact matching step cannot detect non-unique regions shorter than the minimum match length parameter at the beginning or end of candidate ORFs, these regions are removed from the unique regions by subtracting their overlap with the coding sequences of reference transcripts using bedtools subtract (22). Additionally, snoRNA, snRNA, scaRNA and miRNA as well as mt-rRNA and -tRNA, rRNA and rRNA-pseudogene overlapping regions are subtracted, as these genes are often located within introns.

Unique regions are calculated in both transcriptomic and genomic coordinates.

#### Step 3: Find target transcripts by alignment

A candidate Split-ORF transcript contains at least two ORFs that align to the same reference transcript of the same gene, which we term the target transcript. In cases where a candidate Split-ORF transcript has valid alignments to multiple target transcripts, the target transcript with the longest alignments is selected. To find the best matching reference transcript, the non-unique regions identified in step 2 of the Split-ORF pipeline v2 are aligned to the protein-coding reference sequences (Fig. 1C, third step). The exact matches obtained with Mummer3 in step 2 were not used because their strict identity requirement limits sensitivity and an alignment-based approach is employed instead. The alignment can be performed with blastp (v2.17.0+) (23), or diamond (v2.1.12) (24). Alignments are filtered for a minimum length of 15 AA, at least 50% query coverage, and for 95% sequence identity. While the blastp alignment was a part of the first Split-ORF pipeline as well, the usage of the non-unique regions and not the entire ORFs is novel to our method. Using the entire ORF may lead to alignments failing the quality thresholds as the unique regions of the ORFs may reduce the alignment score as well as the identity percentage.

To annotate protein domains, PFAM domains are assigned to the candidate Split-ORFs based on their alignment to the reference protein sequences and the corresponding annotation of the reference protein.

#### Pipeline outputs

Two reports are generated to visually summarize the results of the pipeline run: The Split-ORF report, which presents information about the number of identified candidate Split-ORF transcripts, their functional annotation and the positioning of the candidate Split-ORFs, and the Unique Region Report, which summarizes information about the identified unique DNA and protein regions.

Information on the candidate Split-ORF transcripts, the candidate Split-ORF positioning and their target transcripts is provided in a TSV file. The calculated unique regions are available as BED files, which can be used for visualization in genome browsers and as input for the `ribo-cov` module.

### Processing alternatively spliced transcripts with the split-orf-prediction module

Human transcript sequences annotated as “Nonsense-mediated decay” (NMD), “Retained Intron” (RI) and “protein-coding” were downloaded from Ensembl Biomart (v110). The protein-coding sequences were filtered for transcript support level 1 or 2, or for the presence of their intron chain within the RefSeq annotation (vGCF 000001405.40-RS 2023 10) (25). PFAM annotations and coordinates of exons, coding sequences and coordinates of snoRNA, snRNA, scaRNA and miRNA as well as mt-rRNA and -tRNA, rRNA and rRNA-pseudogenes were downloaded from Ensembl Biomart (v110).

### Ribo-seq coverage in unique regions

As many Split-ORF transcripts arise through alternative splicing and intron retention, a substantial fraction of them is unlikely to be translated. This is because RI transcripts are not always exported to the cytoplasm, and NMD transcripts are rapidly degraded. However, several exceptions have been reported (16; 26; 27). The candidate Split-ORFs are predicted based solely on transcript sequences and therefore need additional evidence to be considered interesting candidates. We examined Ribo-seq coverage in the unique regions of the candidate Split-ORFs. Unique DNA regions were considered supported if they showed significantly higher Ribo-seq coverage than a background set of untranslated regions (UTRs), and were covered by at least three Ribo-seq reads.

### Ribo-seq datasets

We used Ribo-seq data and examined Ribo-seq coverage in the unique Split-ORF regions. The data of Boehm et al. (28) consist of HCT116 cells subjected to UPF1 depletion using degron tags resulting in the inhibition of the NMD pathway. In addition, riboseq.org was queried for Ribo-seq datasets of human cancer cell lines. Two datasets with more than nine samples, treated with cycloheximide, and for which the publication was available were selected. The dataset of Vaklavas et al. (29) containing breast cancer, fibrocystic disease and control samples, referred to as the “breast cancer dataset” from here on, and the glioblastoma dataset of Choudhary et al. (30) containing U343 and U251 cell lines, were obtained with this procedure.

The data of Boehm et al. were downloaded from the BioStudies FTP server (accession E-MTAB-13837). The data of Vaklavas et al. and Choudhary et al. were downloaded from the Sequence Read Archive (SRA) under accession numbers PRJNA523167 and PRJNA591767, respectively.

### Ribo-seq data preprocessing

Preprocessing of the breast cancer dataset of Vaklavas et al. and the glioblastoma dataset of Choudhary et al. was performed by removing the adapter sequences with cutadapt v5.0 (31) and fastp v0.24.0 (32) was used for additional preprocessing and filtering. The Ribo-seq data of Boehm et al. had already been preprocessed. Details about the preprocessing procedure are provided in the Supplementary Materials (Ribo-seq data preprocessing and mapping parameters).

All Ribo-seq data were mapped to the human genome (GRCh38.p14) using STAR (v2.7.11b) (33) with the Ensembl transcriptome annotation (v110). Genomic coordinates of 3’ UTRs and coding sequences were downloaded from Ensembl Biomart (v110) for all protein-coding transcripts and filtered for transcript support level 1 or 2 or for the presence of their intron chain in the RefSeq annotation (vGCF 000001405.40-RS 2023 10).

The Ribo-seq data of Boehm et al. were deduplicated after mapping using UMI tools with default parameters (v1.1.6) (34). Samples of the Vaklavas et al. and Choudhary et al. datasets with fewer than 5 million reads mapping uniquely to annotated genes were excluded from the analysis. RiboseQC (v1.1) (35) was used to assess quality metrics such as 3 nucleotide periodicity of the Ribo-seq samples.

### The ribo-cov module

Assessing the significance of Ribo-seq coverage in the unique DNA regions identified by the Split-ORF pipeline requires a carefully designed statistical framework. These unique regions are expected to show lower translation levels than the non-unique protein-coding sequences, which are shared with the canonical isoforms and are therefore generally translated at higher levels. We performed statistical testing against a background of untranslated regions while taking the length of the unique regions into account.

The steps of the `ribo-cov` module are illustrated in Fig. 4A. Gene expression levels were quantified in transcripts per million (TPM) using htseq-count (v2.0.9) (36) and rnanorm (v2.2.0) (37). The unique regions, coding sequences and the 3’ UTRs were filtered for “translated” genes, by retaining only those genes with a minimum TPM of one.

Furthermore, 3’ UTR regions overlapping coding sequences of other transcript isoforms were removed using bedtools subtract v2.31.1 (22). To estimate background levels, Ribo-seq coverage was quantified in 3’ UTRs and coding sequences in non-overlapping windows of 20 bp (bedtools makewindows `-w 20 -i srcwinnum` and bedtools coverage `-F 0.33 -split`, v2.31.1). An empirical distribution of coding sequence coverage was calculated from all 20 bp windows with at least one mapped read. This empirical distribution represents the coverage distribution of translated 20 bp windows. 3’ UTR windows exceeding the first quartile of the coding sequence distribution were classified as translated and filtered out. The remaining 3’ UTR windows were merged to obtain consecutive untranslated background regions (bedtools merge `-s -c 4,5,6 -o c ollapse,min,distinct`, v2.31.1).

Random regions with the same length distribution as the unique regions were sampled without replacement from the filtered and merged 3’ UTR regions. This procedure was repeated twenty times. For these random regions, length-normalized Ribo-seq coverage was calculated in reads per base (bedtools intersect `-s -wao -sorted` and custom R scripts v4.1.2). The rank-based average of these twenty random 3’ UTR distributions was then calculated, resulting in one background distribution of the Ribo-seq coverage.

Reads in unique regions were considered only if they overlapped the respective region by at least 10 bp. Ribo-seq coverage of the unique regions was calculated in reads per base, and the p-value was obtained with the following formula:

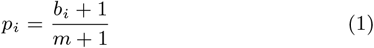

where *b*_*i*_ is the number of random regions exceeding the coverage of unique region i, m is the total number of random regions and *p*_*i*_ is the p-value of unique region i (38). Thus, the p-values correspond to the complementary quantile of the respective unique region within the empirical background distribution. The obtained p-values were adjusted using the Benjamini-Hochberg procedure. Unique regions with a q-value below the FDR threshold of 5% and at least three mapped Ribo-seq reads were considered as significantly Ribo-seq covered (ribo-cov).

The filtering of 3’ UTRs and unique regions, the calculation of the background distribution and of the p- and q-values were performed separately for each sample, resulting in a set of ribo-cov unique regions per sample. These steps are implemented in the ribo-cov module of the `split-orf conda` package. The `ribo-cov` module takes preprocessed FASTQ or BAM Ribo-seq files together with the output of the `split-orf-prediction` step as input. Further details on the implementation, requirements, inputs and parameters of the `split-orf conda` package are provided on GitHub. The `ribo-cov` module provides one CSV file per sample containing the ribo-cov unique regions with their respective q-values and a PDF report summarizing the ribo-cov unique regions across all samples together with plots and result files of the categorization of the candidate Split-ORFs by unique regions and ribo-cov status explained in the following section.

For our analysis, ribo-cov unique regions were further aggregated within the HCT116 NMD inhibited samples, HCT116 control and the cancer samples (breast cancer and glioblastoma datasets) if the unique regions showed significant Ribo-seq coverage in at least two samples. These steps were conducted using custom python scripts and are integrated into the `ribo-cov` module.

### Candidate Split-ORF categorization by unique regions and ribo-cov status

In order to obtain a final set of prioritized candidate Split-ORF transcripts, all candidates were classified based on the presence and position of their unique regions as well as their ribo-cov status. This categorization was used for the sunburst plots shown in Fig. 4B,C created with plotly (39). The pie charts and sunburst plots are automatically generated as part of the `ribo-cov` module.

First, candidate Split-ORF transcripts were divided into ribo-cov analyzable and not ribo-cov analyzable transcripts based on the Ribo-seq coverage of their respective genes and the presence of unique regions within the transcript. A transcript of a gene with a Ribo-seq coverage level above TPM one and the presence of at least one candidate Split-ORF with a unique region was considered ribo-cov analyzable. If the coverage level was below this cut-off, the transcript was assigned to the “not ribo-cov gene” category. Transcripts without unique regions were classified as “no UR”.

For ribo-cov analyzable candidate Split-ORF transcripts, further categorization was based on the unique regions that they contain and the candidate Split-ORFs in which these regions are located. Fig. 3A illustrates the definition of first, middle and last candidate Split-ORFs, referred to as first, middle and last ORFs for simplification. The first ORF of a Split-ORF transcript is defined as the most 5’ ORF, the last ORF as the most 3’ ORF, and any additional ORFs as middle ORFs. For the ribo-cov analyzable classification, the middle and last ORFs were combined into a single category termed “second ORF”, while the “first ORF” retained its definition as the most 5’ ORF. Candidate Split-ORF transcripts were further divided into groups containing unique regions only in the first ORF (“UR 1. ORF”), unique regions only in the second ORF(s) (“UR 2. ORF”) or unique regions in first and second ORFs (“UR 1. & 2. ORF”).

Finally, classification based on the ribo-cov status of the unique regions was performed. The ribo-cov categories were: no coverage in unique regions (“no cov”), ribo-cov in the first ORF (“1. ORF”), ribo-cov in the second ORF (“2. ORF”), or ribo-cov in first and second ORFs (“1. & 2. ORF”). In case of overlapping unique regions (> 20% relative to the shorter unique region) between ORFs of the same transcript, the unique region was assigned only to the more 5’ ORF. For example, if a Split-ORF transcript contains unique regions in the both the first and last ORF that overlap substantially (> 20%), the transcript is classified as “UR 1. ORF” and can only be considered ribo-cov in the first ORF (“1. ORF”).

### GO analyses

GO analyses of the predicted and ribo-cov candidate Split-ORF transcripts was performed using the *g:Profiler* web server (40). For the predicted candidate Split-ORF transcripts, the background set consisted of all genes giving rise to NMD or RI transcripts. For the ribo-cov candidate Split-ORF transcripts, the background was adjusted to all genes giving rise to predicted candidate Split-ORF transcripts with at least one unique region, since only transcripts with unique regions can be evaluated with the `ribo-cov` module.

Only annotated genes were considered for the background set. The displayed terms were manually selected from significantly enriched GO:BP, GO:MF and GO:CC categories using a g:SCS corrected p-value threshold of 5%.

### RNA-binding protein identification

The RBPbase (41) database was downloaded and genes were classified as RBPs if the `any_Hs` was set to `YES` indicating that the gene was identified as an RBP in at least one RNA interactome capture experiment on human data. In total, 4279 genes fulfilled this criterion in RBPbase and were considered as RBPs for the remainder of the analysis.

### Evaluation against the previous pipeline version

To our knowledge, no method predicts Split-ORFs except for the original Split-ORF pipeline. A direct comparison against state-of-the-art tools is therefore not possible. Instead, we evaluated the `split-orf-prediction` module of the Split-ORF pipeline v2 against the Split-ORF pipeline v1. The `ribo-cov` module was not part of the previous pipeline version as it was newly introduced into the extended pipeline.

## Results

We define a Split-ORF transcript as one that contains two or more ORFs that encode parts of the same full length protein, as illustrated in Fig. 1A. The Split-ORF pipeline predicts candidate Split-ORF transcripts based on a transcript’s sequence. For this prediction, the possible ORFs within each transcript are identified, their match to the original protein-coding reference transcript (target) is determined by sequence alignment, and unique regions that are not present in the target transcript are calculated. These steps are illustrated in Fig. 1C. According to our definition, a transcript is considered a candidate Split-ORF transcript if it contains two high-quality alignments to the same target protein coding sequence.

### Prediction of candidate Split-ORFs from alternatively spliced transcripts

#### The Split-ORF pipeline v2 predicts thousands of candidate Split-ORF transcripts

Split-ORFs can arise through the skipping or inclusion of alternative exons that change the reading frame and thereby introduce premature termination codons. They can also arise through the inclusion of poison cassette exons or the retention of introns. To identify all candidate Split-ORF transcripts the extended Split-ORF pipeline’s `split-orf-prediction` module was applied to human transcripts annotated as RI and NMD in Ensembl (v110). Strikingly, 43.0% (9,185) of the NMD transcripts were predicted by our pipeline to have the potential to form Split-ORFs, although only 4,435 unique DNA regions were detected (Table 2). In contrast, only 16.2% (5,530) of the RI transcripts were predicted as candidate Split-ORF transcripts with 7,219 unique DNA regions identified (Table 2).

**Table 2.** Split-ORF prediction and ribo-cov results for nonsense-mediated decay (NMD) and retained intron (RI) transcripts (Ensembl v110). SO: candidate Split-ORF, UR: unique region, ribo-cov: significant Ribo-seq coverage compared to untranslated background. The set of ribo-cov transcripts and URs exclude cases of significant Ribo-seq coverage in the first candidate Split-ORF of the NMD inhibited HCT116 samples; for the SOs column, all ribo-cov SOs are counted, including the ribo-cov first SOs in NMD inhibition.

| Type | Genes | Transcripts | SOs | URs |
| --- | --- | --- | --- | --- |
| Ensembl NMD | 8342 | 21384 | / | / |
| Split-ORF trans | 4119 | 9185 | 19802 | 4435 |
| ribo-cov | 278 | 433 | 781 | 453 |
| Ensembl RI | 11291 | 34128 | / | / |
| Split-ORF trans | 2558 | 5530 | 12901 | 7219 |
| ribo-cov | 524 | 1062 | 1367 | 1174 |

We found that the majority of candidate Split-ORF transcripts contain two ORFs, while a substantial fraction also contains three ORFs. Candidate Split-ORF transcripts with four or more ORFs are rare, as shown in Fig. 2B. The mean Split-ORF lengths are 658 bp and 873 bp for RI and NMD transcripts, respectively. Most candidate Split-ORFs are shorter than 1 kilobase (kb), and only a minor fraction exceeds 2 kb (Fig. 2A).

**Figure 2.**
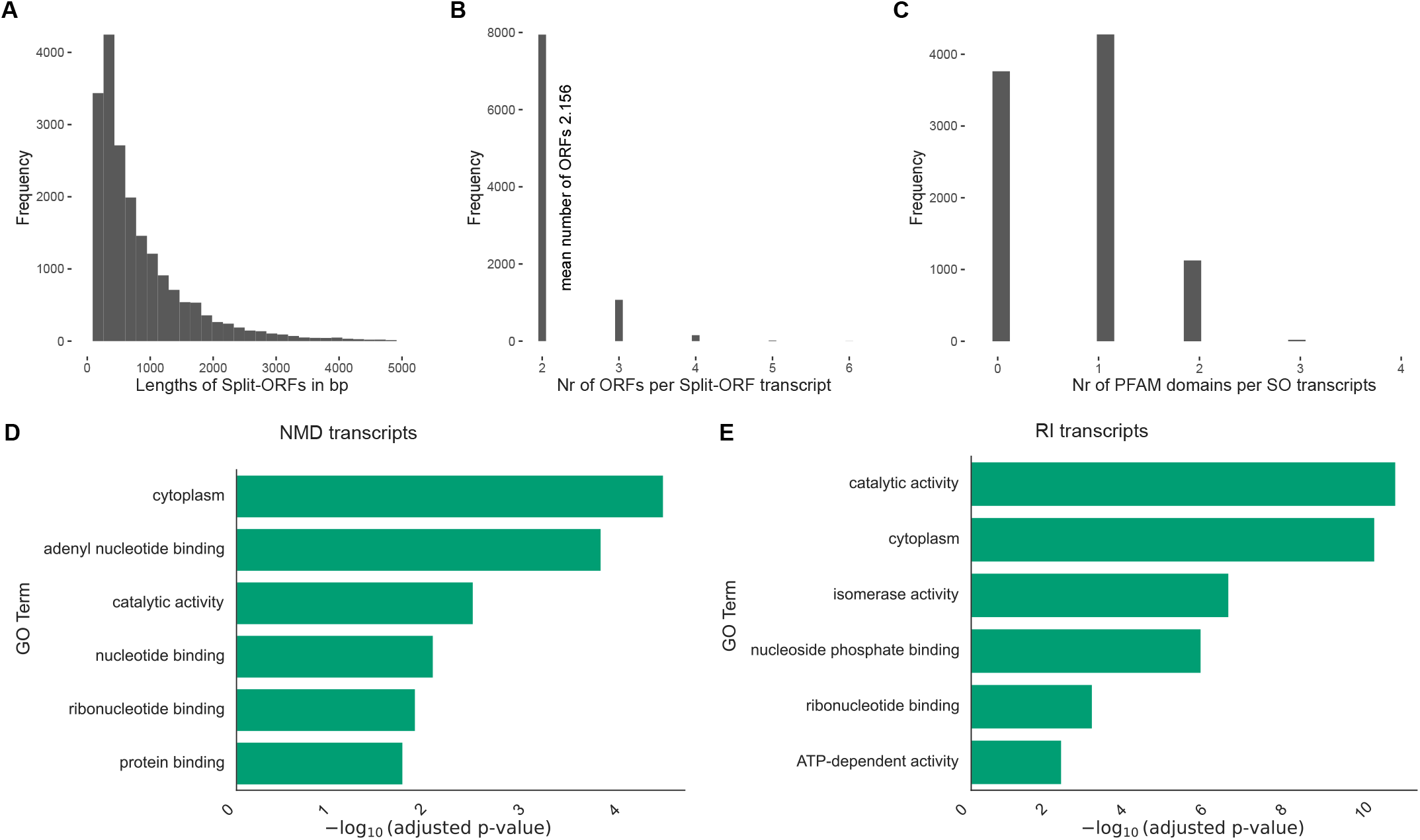
Predicted candidate Split-ORFs are enriched for nucleotide binding terms. **A**: Histogram of the length of the candidate nonsense-mediated decay (NMD) Split-ORFs in bp. **B**: Histogram of the number of candidate Split-ORFs per NMD candidate Split-ORF transcript. **C**: Histogram of the number of PFAM domains overlapping NMD candidate Split-ORF (SO) transcripts. **D**: GO enrichment of the genes giving rise to NMD candidate Split-ORF transcripts. **E**: GO enrichment of the genes giving rise to retained intron (RI) candidate Split-ORF transcripts.

To test whether candidate Split-ORFs occur in particular protein domains, we used annotations from the PFAM database. Indeed, 2520 RI and 5422 NMD candidate Split-ORF transcripts overlap with one or more annotated protein domains. As illustrated in Fig. 2C, more than half of the NMD candidate Split-ORF transcripts overlap with at least one PFAM domain. In contrast, among RI transcripts, 1910 overlap with one and 610 transcripts overlap with 2 PFAM domains. The most common PFAM domain of both transcript types is PF00400, a WD40 domain which is frequently involved in protein-protein interactions and complex assembly.

GO term enrichment analysis of the genes giving rise to the candidate Split-ORF transcripts (4119 NMD and 2558 RI genes) revealed that nucleotide binding terms are highly enriched in both groups (Fig. 2D,E). In addition, protein binding is enriched among the NMD transcripts, whereas RI transcripts are enriched for catalytic and isomerase activities.

#### The positioning of unique regions is consistent with poison cassette exons and RIs driving Split-ORF formation

For both NMD and RI candidate Split-ORF transcripts, we identified 4435 and 7219 unique regions, respectively, enabling the large scale testing for significant Ribo-seq coverage (Table 2). Among NMD candidate Split-ORF transcripts, 5800 contain no unique region, 2564 contain one unique region and 821 contain two or more unique regions (Fig. 3B). Because unique regions of different candidate Split-ORFs may overlap, applying a maximum overlap threshold of 20% reduced the number of NMD candidate Split-ORF transcripts with two unique regions to 522. The majority of RI candidate Split-ORF transcripts harbors one unique region (2521), and 1686 RI candidate Split-ORF transcripts contain two or more unique regions. 1506 RI candidate Split-ORF transcripts fulfill the criterion of having two or more unique regions that do not overlap more than 20%.

**Figure 3.**
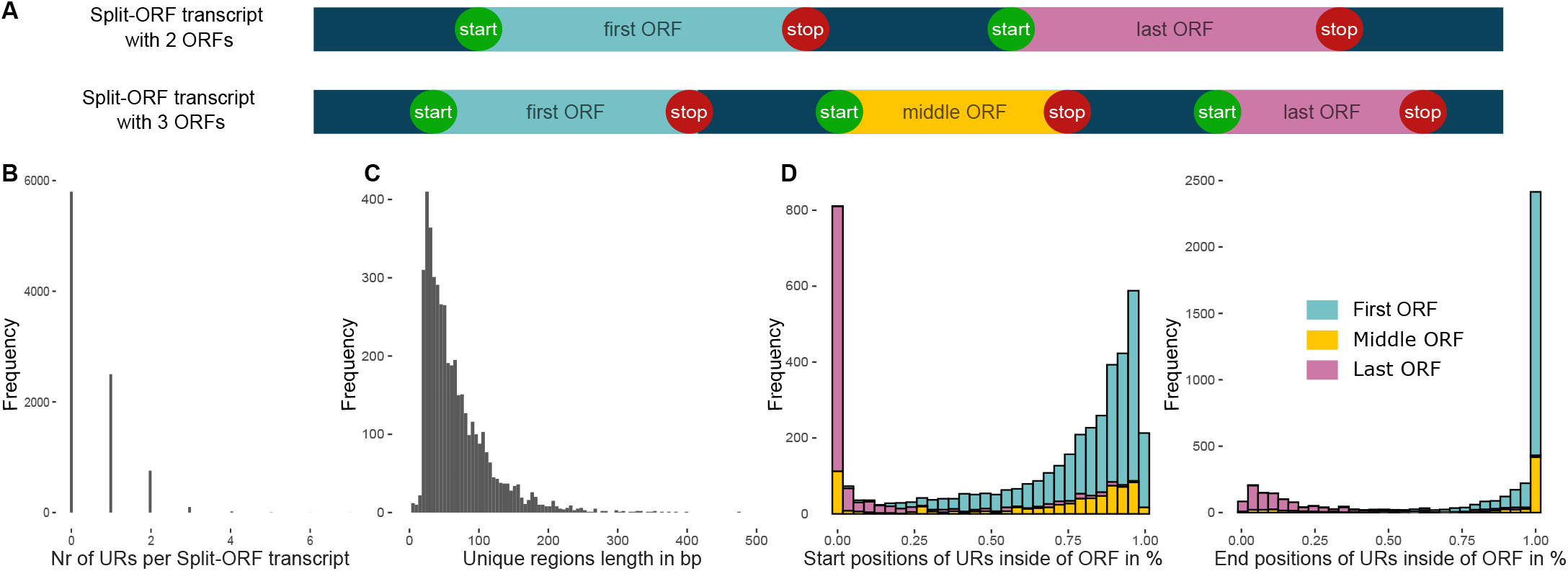
Unique Split-ORF regions are positioned at the end of the first and the beginning of the last ORF. **A**: The first ORF is the most 5’ ORF in a Split-ORF transcript, the last ORF is the most 3’ ORF in a Split-ORF transcript. If more than 2 ORFs exist, there are additional middle ORFs located between the first and the last ORF. **B**: Histogram of the number of unique DNA regions per NMD candidate Split-ORF transcript. **C**: Histogram of unique DNA regions’ length in bp for NMD candidate Split-ORF transcripts. **D**: Left: Histogram of the start positions of the unique regions in % with respect to the candidate Split-ORF. Right: Histogram of the end positions of the unique regions in % with respect to the candidate Split-ORF. Both plots are colored by ORF position in the candidate Split-ORF transcript.

Unique regions are usually short, with median lengths of 53 and 75 bp for NMD and RI candidate Split-ORF transcripts, respectively, as illustrated in Fig. 3C. Only a few unique regions are longer than 1 kb, with the largest spanning 2446 bp.

As Split-ORF transcripts have two or more Split-ORFs, we defined the first ORF as the most 5’ Split-ORF and the last ORF as the most 3’ Split-ORF (Fig. 3A). Split-ORF transcripts with three or more ORFs also contain one or several middle ORFs which are located between the first and the last ORF. Fig. 3D shows the positions of the starts and ends of the unique regions with respect to the different candidate Split-ORFs (first, middle and last). For the first ORFs, the unique regions tend to end at the stop codon of the ORFs and rarely in the middle of the ORFs (Fig. 3C, right panel). In contrast, unique regions in the last ORFs, tend to start at the beginning of the ORFs and rarely in the middle (Fig. 3C, left panel). These positions are consistent with a scenario in which a poison cassette exon or a RI introduces a premature termination codon, creating the first Split-ORF, while also introducing a new in-frame start codon that gives rise to the second Split-ORF, as reported for SRSF7 (16) and as illustrated in Fig. 1A and B. The supplemental information contains an example of a unique region ending in the middle of a candidate Split-ORF, to illustrate why these regions might be informative and should not be filtered out.

### Evaluation of Split-ORF prediction against the previous pipeline version

The first version of the Split-ORF pipeline was run on the same NMD and RI transcripts and resulted in 8981 NMD candidate Split-ORF transcripts and 5509 RI candidate Split-ORF transcripts. Compared to the predictions of the Split-ORF pipeline v2 (Table 2), these are 204 and 21 less candidate Split-ORF transcripts for NMD and RI, respectively. The increase in the number of candidate Split-ORF transcripts with the new pipeline can be attributed to the use of the non-unique regions for the alignment step instead of the entire ORF.

### Ribo-seq coverage in unique regions

The Split-ORF pipeline v2 was augmented by the `ribo-cov` module, because such a functionality was missing. The previous version relied on a manual procedure of checking whether poison cassette exons contained at least two Ribo-seq reads. The new `newribo-cov` module performs statistical testing of coverage in unique regions against an untranslated background. A comparison of the `ribo-cov` functionality between pipeline versions is therefore not possible.

The `ribo-cov` module calculates Ribo-seq coverage in the unique regions and 3’ UTRs. From the 3’ UTR coverage, a background distribution is inferred. Unique regions are then tested for significantly higher Ribo-seq coverage relative to this background, as illustrated in Fig. 4A. This results in a set of “ribo-cov” unique regions and is performed at the sample level. Further categorization of the candidate Split-ORF transcripts based on which unique regions are ribo-cov is performed as described in the methods; obtaining the categories of the first (most 5’) Split-ORF, the second (middle or most 3’) Split-ORF or first and second Split-ORFs being ribo-cov. These are referred to as first and second ORF.

**Figure 4.**
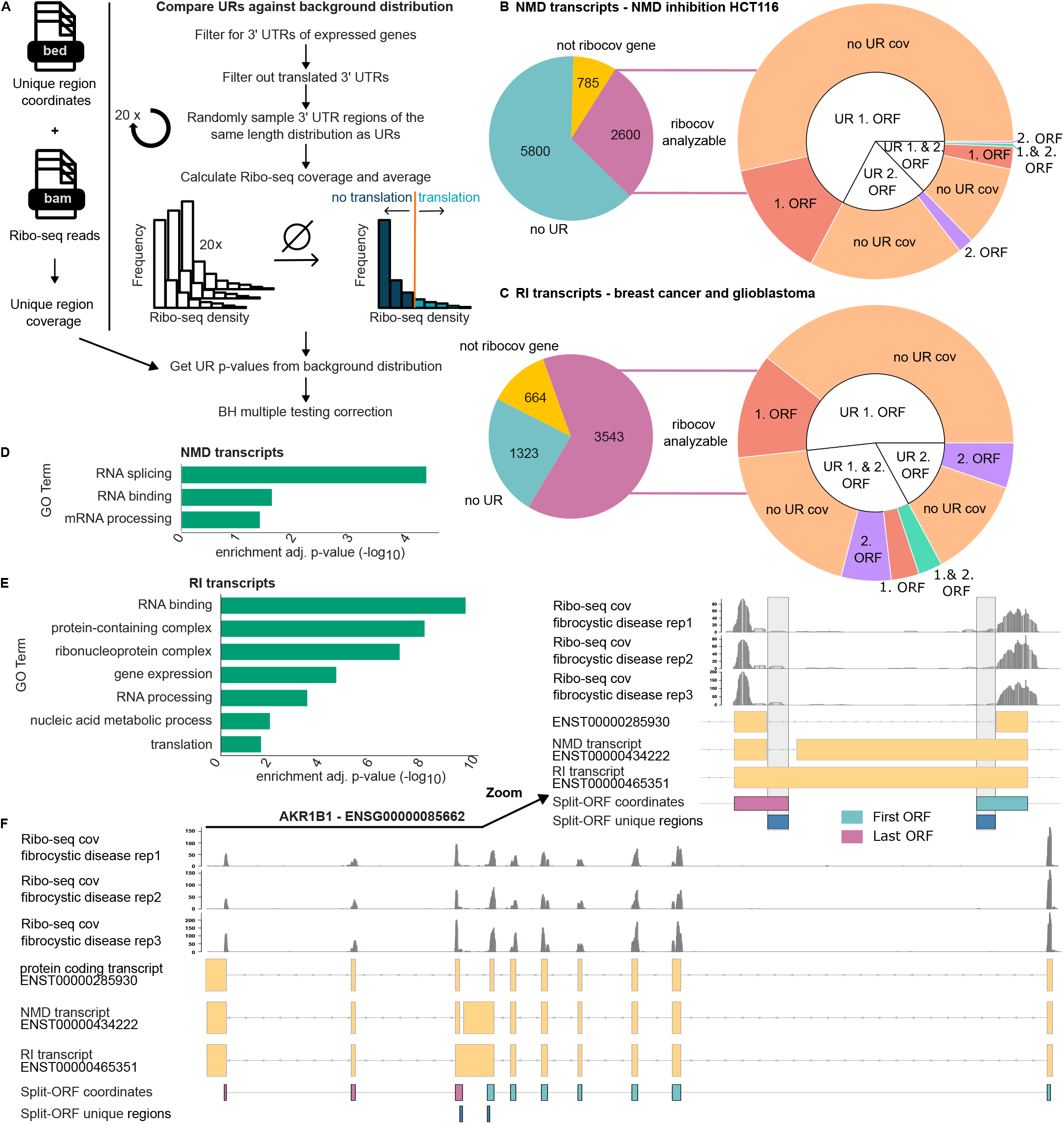
Unique Split-ORF regions with significant Ribo-seq coverage in diverse cell types and diseases. **A**: Ribo-seq coverage in unique regions (URs) is tested against a background distribution of 3’ untranslated regions (UTRs). URs with significantly higher coverage are termed ribo-cov. **B**: Left: Pie chart indicating the number of ribo-cov analyzable NMD candidate Split-ORF transcripts with the HCT116 NMD inhibited samples. (Right) Sunburst chart of ribo-cov NMD candidate Split-ORF transcripts by unique region position for the HCT116 NMD inhibited samples. **C**: Left: Pie chart indicating the number of ribo-cov analyzable RI candidate Split-ORF transcripts with the glioblastoma and breast cancer samples. (Right) Sunburst chart of ribo-cov RI candidate Split-ORF transcripts by unique region position for the glioblastoma and breast cancer samples. **D**: GO enrichment of interesting candidate Split-ORF genes of the NMD transcripts. **E**: GO enrichment of interesting candidate Split-ORF genes of the RI transcripts. **F**: Example of Ribo-seq coverage in *AKR1B1* transcripts and unique regions of RI candidate Split-ORF transcript ENST00000465351 in fibrocystic disease samples. The y-axis of the Ribo-seq coverage is given in read depth per base.

Publicly available Ribo-seq data of HCT116 cells with and without NMD inhibition, as well as breast cancer and glioblastoma cell lines were used for this analysis. Information on the samples, including mapping rates and read counts is provided in Table S1.

#### Hundreds of candidate Split-ORF transcripts showed significant Ribo-seq coverage

The interpretation of ribo-cov unique regions depends on the biological condition of the samples. While NMD transcripts are normally degraded, inhibiting the NMD pathway in HCT116 cells may result in their accumulation and in the translation of truncated protein products. If the first ORF of a candidate Split-ORF transcript is translated under NMD inhibition, this does not constitute evidence for Split-ORF formation. In contrast, if the first and second ORFs, or only the second ORF is translated, this observation cannot be explained by NMD inhibition alone. For cancer and HCT116 control samples, however, NMD transcripts are expected to be rapidly degraded. Observing ribo-cov first ORFs would be consistent with our previous hypothesis that translation of the second ORF stabilizes the transcript and thereby promotes the translation of the first ORF. Therefore, we define interesting Split-ORF candidate transcripts and genes as being ribo-cov in any Split-ORF in the cancer and HCT116 control samples, or being ribo-cov in at least a second ORF for the NMD inhibited HCT116 samples.

Table 3 summarizes the ribo-cov candidate Split-ORFs for the union of all cancer samples, the HCT116 control samples, as well as the HCT116 NMD inhibited samples. Comparing the numbers of ribo-cov NMD transcripts in the NMD inhibited HCT116 samples (Fig. 4B) with the HCT116 control samples, the amounts of ribo-cov first, second as well as first and second ORFs all increase. For the HCT116 control samples, ribo-cov unique regions are rare, 26 first ORFs are ribo-cov and only two second ORFs show significant Ribo-seq coverage (Table 3). As the increase in the ribo-cov first ORFs can be attributed to the translation of a “faulty” truncated protein under NMD inhibition, these cases were not considered further for the NMD inhibited samples. Hence, most ribo-cov candidate Split-ORFs considered in this analysis are the ribo-cov first ORFs of the glioblastoma and breast cancer samples. Ten NMD candidate Split-ORF transcripts are ribo-cov in the first and second ORF upon NMD inhibition, whereas none of the candidate Split-ORF transcripts are ribo-cov in both ORFs in the HCT116 control samples (Table 3). This finding suggests that some NMD transcripts encode Split-ORFs and may be translated if they are stabilized.

**Table 3.**
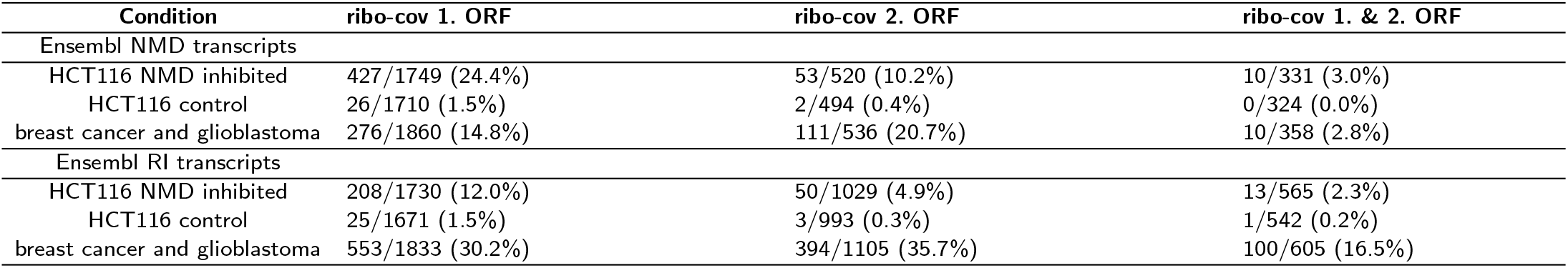
Ribo-cov transcripts by unique region positions. The number of transcripts with ribo-cov unique regions in the first, second or first and second ORFs are given with respect to the number of transcripts that contained unique regions in the first, second or first and second ORFs, respectively. Candidate Split-ORF transcripts were considered if their corresponding genes were detected in at least two samples with a TPM of at least one.

| Condition | ribo-cov 1. ORF | ribo-cov 2. ORF | ribo-cov 1. & 2. ORF |
| --- | --- | --- | --- |
| Ensembl NMD transcripts |  |  |  |
| HCT116 NMD inhibited | 427/1749 (24.4%) | 53/520 (10.2%) | 10/331 (3.0%) |
| HCT116 control | 26/1710 (1.5%) | 2/494 (0.4%) | 0/324 (0.0%) |
| breast cancer and glioblastoma | 276/1860 (14.8%) | 111/536 (20.7%) | 10/358 (2.8%) |
| Ensembl RI transcripts |  |  |  |
| HCT116 NMD inhibited | 208/1730 (12.0%) | 50/1029 (4.9%) | 13/565 (2.3%) |
| HCT116 control | 25/1671 (1.5%) | 3/993 (0.3%) | 1/542 (0.2%) |
| breast cancer and glioblastoma | 553/1833 (30.2%) | 394/1105 (35.7%) | 100/605 (16.5%) |

The numbers in Table 3 are based on the transcript level. As different NMD and RI isoforms may arise from the same gene, it is also informative to consider the number of genes giving rise to the ribo-cov Split-ORF candidates. For example the ten NMD transcripts that are ribo-cov in both the first and second ORF in the HCT116 NMD inhibited samples belong to eight different genes, namely: *SREBF2, BCLAF1, GLOD4, C1orf174, DHX35, STAT2, ALG5* and *TAF2*. Supplementary Figure 2 shows three examples (*BCLAF1, DHX35* and *TAF2*) of NMD candidate Split-ORF transcripts which are ribo-cov in both unique regions.

The largest number of candidate Split-ORF transcripts with coverage in both first and second ORFs was found among RI transcripts in the glioblastoma and breast cancer samples, namely 100 candidate Split-ORF transcripts. These 100 transcripts arise from 61 different genes, including *DDX24, DDX5, AKR1B1* and *PABPC4* to name some examples. Fig. 4F illustrates an RI transcript of *AKR1B1* and the corresponding Ribo-seq coverage in the unique regions in the fibrocystic disease samples.

Taken together, there are 48 interesting candidate Split-ORF genes for the NMD transcripts in the HCT116 NMD inhibited samples, 18 for the HCT116 control samples and 253 for NMD transcripts in the glioblastoma and breast cancer samples, 33 for RI transcripts in the HCT116 samples with NMD inhibition, 13 in the HCT116 control samples and 516 interesting candidate genes for the RI transcripts in the glioblastoma and breast cancer samples. Taking the union of these genes for the NMD and RI transcripts, resulted in 278 and 524 interesting candidate genes, respectively (Table 2).

#### The majority of the interesting candidate Split-ORF genes encodes RNA-binding proteins

For the 278 NMD genes and the 524 RI genes that gave rise to candidate Split-ORF transcripts prioritized by the `ribo-cov` module, a GO term analysis was performed. RNA-binding and RNA-processing terms were enriched for both NMD and RI transcripts (Fig. 4D, E). Compared to the GO enrichment of all candidate Split-ORF genes in Fig. 2D,E, where more general binding terms such as nucleotide, ribonucleotide and protein binding were enriched, RNA-binding and RNA-processing functions emerged in our ribo-cov candidate Split-ORF genes. This suggests that a substantial fraction of interesting candidate Split-ORF genes encodes RBPs.

To further investigate whether the ribo-cov Split-ORF candidates encode RBPs, RBPbase was queried. Of the 278 interesting NMD candidate Split-ORF genes, 134 are classified as RBPs according to RBPbase. Furthermore, 301 of the 524 RI candidate Split-ORF genes are annotated as RBPs. This strengthens our hypothesis that Split-ORF formation is a shared mechanism among different RBPs.

## Discussion

Split-ORFs occur on transcripts with two or more ORFs, each of which encodes a part of the same full length protein. We extended and improved the Split-ORF pipeline, which predicts the ability of transcripts to encode Split-ORFs based on their nucleotide sequence. Applying the split-orf-prediction module to human RI and NMD transcripts resulted in the prediction of more than 14,000 candidate Split-ORF transcripts. While these predictions represent only the potential of transcripts to encode Split-ORFs, the pipeline also identified more than 11,000 unique regions that enable to detect Split-ORF specific signals in Ribo-seq data.

The extended Split-ORF pipeline introduces unique regions and performs statistical testing of Ribo-seq coverage via the new `ribo-cov` module, which had no counterpart in the first version. Candidate Split-ORF transcripts arising from splicing alterations other than poison cassette exons can now also be tested, and the procedure applies directly to RI transcripts without manual intervention. The Split-ORF pipeline v2 can be applied directly to different Ribo-seq data sets, organisms, custom transcriptome assemblies or other sets of alternatively spliced transcripts. The testing procedure of the Ribo-seq coverage in the unique regions against an untranslated background, provides p- and q-values, enabling candidates to be assessed on a statistical basis. Information on which ORF of the transcript shows the Ribo-seq coverage (first or second ORF) is reported and an assignment into different confidence categories based on the unique region position and ribo-cov status of the transcripts is automatically obtained.

In the `split-orf-prediction` module, using the non-unique regions for the alignment step identified more candidate Split-ORFs than using the entire ORF, a smaller but additional improvement. For both modules, the pipeline now summarizes the results in the form of HTML or PDF reports.

A higher number of NMD candidate Split-ORF transcripts was predicted, but unique regions were more frequent among RI transcripts. NMD transcripts may form Split-ORFs by different mechanisms. In addition to the inclusion of poison cassette exons, exon skipping or alternative splice donor and acceptor sites may introduce premature termination codons into the ORFs. While alternative splice sites and exon skipping do not necessarily introduce unique sequences into the Split-ORFs, RIs always incorporate novel sequences into the transcript, resulting more often in one or more unique Split-ORF regions per transcript. The absence of unique regions in candidate Split-ORFs is a major limitation for identifying interesting Split-ORF candidates. For the NMD transcripts only ∼25% of the candidate Split-ORFs contain a unique region and the majority of RI candidate Split-ORF transcripts contains no or only a single unique region. Transcripts in which both candidate Split-ORFs contain a unique region are less frequent and only a fraction of them showed ribo-cov in both unique regions. The strength of the `ribo-cov` module’s testing procedure is that it reduces the large set of candidate Split-ORFs to a manageable set of candidate transcripts and genes for further targeted investigation.

Using ribo-cov in the first candidate Split-ORF as evidence for Split-ORF formation is based on the hypothesis that translation of the second Split-ORF removes all bound exon-junction complexes and thereby protects the transcript from NMD in cis. While this was demonstrated for SRSF7 (16), it remains to be determined whether this mechanism applies to other Split-ORFs. Cases in which only the second ORF is ribo-cov may arise when no unique region is present in the first ORF. If both first and second ORFs have unique regions, but only the second ORF is ribo-cov, it is possible that the pipeline predicted the first ORF incorrectly. Alternatively, another unknown mechanism may promote the translation of the second ORF. Consequently, the Split-ORF candidates with ribo-cov in only one unique region should be considered weaker evidence for Split-ORF formation than the candidates in which both unique regions are ribo-cov.

When dealing with different candidate Split-ORF isoforms of the same gene, unique regions may overlap or be identical. For identical unique regions, it is not possible to know from which transcript the observed Ribo-seq coverage comes, it may also be the case that both transcript isoforms are ribo-cov. Figure S2A and C show *BCLAF1* and *TAF2* candidate Split-ORF transcripts whose unique regions overlap. For *BCLAF1* the second unique region of transcript ENST00000534269 is identical to the first unique region of transcript ENST00000527613. Hence, it cannot be determined whether both candidate Split-ORFs of transcript ENST00000534269 are ribo-cov, or whether the translation of the first ORFs of both transcripts gives rise to the observed profile. In general, isoforms missing from the reference protein-coding transcripts may also overlap with the unique regions. While we used the Ensembl annotated protein-coding transcripts with transcript support level 1 or 2 or the presence of their intron chain within the RefSeq annotation, the reference set of transcripts can be configured by the user to match the requirements of a given analysis.

Furthermore, observing significant Ribo-seq coverage in the candidate Split-ORF unique regions suggests translation, but does not constitute definite proof. Mapping artifacts, binding of RBPs, or secondary RNA structures may cause sequence fragments of similar length as ribosomal footprints. We therefore designed a background of untranslated regions to rigorously test unique region coverage, balancing the risk of calling spurious ribosomal footprint mappings as significant, while avoiding being too conservative as Split-ORF unique regions may be translated at much lower levels than canonical coding sequences. Despite this effort, false positive significant candidate Split-ORF unique regions cannot be precluded. We, therefore, consider the ribo-cov unique regions not as conclusive proof for Split-ORF translation, but as strong clues which are promising targets for validation in future functional studies.

The enrichment of nucleotide-binding terms among all candidate Split-ORF transcripts, and RNA-binding and RNA-processing terms among ribo-cov genes is in line with our previous findings (16), where we proposed Split-ORF formation as a possible mechanism shared among many RBPs. The enrichment of other terms among all candidate Split-ORF transcripts, which are absent from the ribo-cov-supported candidates, may indicate false positive predictions that do not form translated Split-ORFs. For example, some RI transcripts may be kept in the nucleus and are not translated at all (42).

The NMD and RI transcripts used in this study are based on general annotations and lack cell type specificity. Because splicing is highly cell type-specific (43; 44; 45), some annotated transcripts may not be expressed in the cell types analyzed here, while the analyzed cell types may have additional Split-ORF transcripts which were missed here as they are not represented in the annotation.

## Limitations

This study introduces the Split-ORF pipeline v2, which predicts candidate Split-ORFs and tests for significant Ribo-seq coverage to reduce the predictions to a set of promising candidates. Investigating the mechanism of Split-ORF formation requires functional experiments which are interesting future directions for the field but beyond the scope of this work.

As discussed above, testing for significant Ribo-seq coverage cannot be considered as conclusive validation of Split-ORF translation. An obvious improvement would be to analyze the three nucleotide periodicity within the unique regions to test whether the correct reading frame is translated. Testing periodicity for the entire Split-ORF will generate a lot of false positives as the non-unique protein-coding sequences are part of the candidate Split-ORF transcripts but of other canonical mRNAs as well. Most tools, however, do not allow for the definition of custom regions to test. One tool that allows to input regions of interest is RiboTISH (46). However, as it was build to test for periodicity in ORFs, the short length of the unique regions results in a large decrease of statistical power rendering the procedure ill-suited for testing Split-ORF candidates (see Github-RiboTISH-issue-47).

Unequivocal validation of Split-ORF proteins would require proteomics datasets which capture peptides unique to the candidate Split-ORF proteins. A unique peptide is distinct from all peptides of canonical proteoforms and the trypsin digestion pattern determines its exact sequence and size. This is a very different type of uniqueness than the unique regions calculated by the Split-ORF pipeline. A unique peptide region from the Split-ORF pipeline may not be detectable by mass spectrometry due to missing or too frequent tryptic cleavage sites, ions which are not ionizable and general issues with sparseness. Furthermore, Wang et al. (47) demonstrated that the tryptic cleavage used for most proteomics datasets limits the detection of alternatively spliced isoforms due to evolutionary conserved preference of lysin and arginine codons at exon boundaries. This prevents the generation of unique peptides at exon-exon junctions. As Split-ORFs occur in alternatively spliced transcripts, validation of their translation via unique peptides in proteomics data following tryptic digestion protocols suggests limited potential for success. We, therefore, decided to focus this study solely on Ribo-seq coverage for Split-ORF corroboration.

## Conclusion

The exact mechanism of Split-ORF translation and function and how cell type specificity takes effect constitute future directions of the field. The Split-ORF pipeline v2 is the first step towards the computationally guided identification and validation of Split-ORFs. For NMD and RI transcripts combined, we identified 715 distinct genes giving rise predicted candidate Split-ORF transcripts which had significant Ribo-seq coverage in at least the second candidate Split-ORF for the NMD inhibited HCT116 samples or any candidate Split-ORF for the glioblastoma, breast cancer and HCT116 control samples. The majority of these candidate genes encode RBPs. The Split-ORF pipeline v2 recovered Split-ORF candidates in different confidence sets well beyond the originally described *SRSF7* which strengthens the hypothesis that Split-ORF formation may be a more wide-spread phenomenon especially prevalent in RBPs.

## Code and Data availability

The code of the Split-ORF pipeline is available online Github, with instructions on how to install the conda package as well as which in- and output files are required. The pipeline for the preprocessing of the Ribo-seq datasets for usage with the ribo-cov module of the Split-ORF pipeline is available on GitHub. The Input files to the Split-ORF pipeline together with all results are deposited on Zenodo.

## Acknowledgements

This work was supported by the Goethe University Frankfurt am Main, the German Centre for Cardiovascular Research (DZHK Standort Rhine Main 81Z0200101 to M.H.S., C.K.), the Deutsche Forschungsgemeinschaft (DFG) excellence cluster EXS2026 (Cardio-Pulmonary Institute, project-ID 390649896 to M.H.S. and C.K. and M.M-M.),), the DFG project-ID 403584255 - TRR 267 (TP Z03 to M.H.S. and TP A03 to M.M-M.). M.H.S. acknowledges the Hessian.AI for funding.

