## Supplementary Information for "The extended Split-ORF pipeline: Prediction and evaluation of Split-ORFs using Ribo-seq data"

### 1 Pipeline implementation

The Split-ORF pipeline is implemented in a modular fashion within the bash framework. The different modules of the pipeline are written in python (v3.11) and the reports are generated with R (v4.1.2) with the Rmarkdown framework. The pipeline is available as a conda package (https://github.com/SchulzLab/SplitOrfs), we are planning to make it available via bioconda.

**Input and output files** In order to run the `split-orf-prediction` module of the Split-ORF pipeline, the following inputs are required: (1) sequences of the transcripts of interest in FASTA format, (2) FASTA files of protein coding reference transcripts and (3) proteins, (4) a TSV file with PFAM annotations, (5) the exon coordinates of the transcripts of interest, (6) the alignment method (blast or diamond), exonic coordinates of contaminating gene sequences (snoRNA, miRNA, snRNA, scaRNA, rRNA, rRNA_psuedogene, Mt_tRNA, Mt_rRNA) and (8) a BED file with the genomic coordinates of the coding sequences (CDSs) of the reference transcripts.

All of these input files were downloaded from Ensembl Biomart (Ensembl Genes 110, GRCh38p.14) using the following mart options and transformed with the respective custom scripts which can be found in the Input_scripts folder of the Split-ORF pipeline GitHub repository: https://github.com/SchulzLab/SplitOrfs.

(1) Attributes: Sequences: cDNA sequences, header information: Gene stable ID, Transcript stable ID, Filters: Gene: Transcript type: nonsense_mediated_decay or retained_intron.

(2) Attributes: Sequences: cDNA sequences, header information: Gene stable ID, Transcript stable ID, Filters: Gene: Transcript type: protein_coding.

(3) Attributes: Sequences: Peptide, header information: Gene stable ID, Transcript stable ID, Filters: Gene: Transcript type: protein_coding.

(4) Attributes: Features: Gene: Gene stable ID Transcript stable ID; Protein Domains and Families: Pfam ID, Pfam start, Pfam end. Remodelling scripts: `convert_ensembl_output_to_bed.py`.

(5) Attributes: Structures: Gene stable ID, Transcript stable ID, Exon region start, Exon region end, Transcript start, Transcript End, Strand. No filters were selected. Downloading all exonic positions includes those of the transcripts of interest.

(7) Attributes: Structures: Chromosome/scaffold name, Gene stable ID, Transcript stable ID, Exon region start, Exon region end, Strand. Filter: Gene: the respective contaminating RNA.

(8) Attributes: Structures: Chromosome/scaffold name, Gene stable ID, Transcript stable ID, Genomic coding start, Genomic coding end, Strand.

Protein coding transcript (2) and protein sequences (3) as well as CDS coordinates (7) were filtered for transcript support level 1 or 2 or the presence of their exact intron chain in the RefSeq annotation (`vGCF_000001405.40-RS_2023_10`, Pruitt et al. 2014) with custom scripts (`filter_Ensembl_GTF.sh`, `Filter_prot_coding_reference.sh`). The filtered CDS coordinates were combined with the contamination coordiantes into a single BED file using the remodelling script: `generate_CDS_contamination_subtraction_coordinates.sh`. The input files are specified via a JSON file and all input data should be located in the same directory. The Split-ORF pipeline creates an output folder with a timestamp of the run at a user specified location. All results as well as intermediate result files are written into this output directory. The final output files are a TSV file of the predicted Split-ORFs, BED files of the genomic coordinates of the unique Split-ORF regions and two HTML reports about the predicted Split-ORF candidates and about their unique regions. The steps of the Split-ORF pipeline produce intermediate results which are also included in the output of the pipeline.

The Input and Output files for the `ribo-cov` module are described in more detail on GitHub.

### 2 Filtering of unique regions

#### 2.1 Keeping unique regions in the middle of Split-ORFs

The proposed model with the poison cassette exon or retained intron introducing unique sequences at the end of the first Split-ORF and the start of the second Split-ORF raises the question of how unique regions can appear in the middle of Split-ORFs and whether these should be kept. One example of how these middle unique regions may arise is if there is a small part of the gene that has an exact sequence match to a different gene within a "unique region". For example, transcript ENST00000412573 of gene ENSG00000164647 has two unique regions for its Split-ORF with ID ORF-2854 (as assigned by the Split-ORF pipeline): namely 218-257 and 278-304 in transcriptomic coordinates. There is a gap of 21 bp between these two unique regions which shares sequence identity with another gene ENSG00000196476 or C20orf96. Both unique regions are hence "valid" unique regions and should be kept.

#### 2.2 Unique regions overlapping CDS coordinates

As stated in the methods section of this paper, overlapping parts of the unique DNA regions with the reference transcripts' CDSs were removed. Unique regions can overlap with CDS coordinates only if a non-unique region is present at the start or end of the respective Split-ORF and this region is smaller than the mummer length parameter of 20 bp. Mummer3 can only find exact matching regions, it cannot explicitly look for regions that do not match exactly. Regions with exact matches smaller than the required length parameter are hence considered as unique. The removal of overlapping parts of the unique DNA regions with the reference transcripts' CDSs remedied the inclusion of these false unique regions.

### 3 Ribo-seq data preprocessing and mapping parameters

Preprocessing of the breast cancer dataset of Vaklavas et al. (Vaklavas et al. 2020) and the glioblastoma dataset of Choudhary et al. (Choudhary et al. 2020) was performed by removing the adapter sequences "AGATCGGAAGAGCACACGTCTGAACTCCAGTCAC" and "CTGTAGGCACCATCAAT", respectively, using cutadapt v5.0 (Martin 2011) with the following parameters `--minimum-length 25`, `--max-n 0.1`, `--max-expected-errors 1` and `-a` followed by the adapter sequence. Fastp v0.24.0 (Chen et al. 2018) was used for additional preprocessing on these datasets with the following parameters `--length_required 25`, `--cut_front`, `--cut_front_window_size 1`, `--cut_mean_quality 15`, `--cut_tail`, `--cut_tail_window_size 1` and `--length_limit 35`.

The Ribo-seq data of Boehm et al. (Boehm et al. 2024) were downloaded as BAM files. The contained reads had already been preprocessed, adapters had been removed and the unique molecular identifiers (UMIs) had been added to the read names. These BAM files were converted to FASTQ files (bedtools bamtofastq v2.31.1, Quinlan et al. 2010).

All Ribo-seq data were mapped to the human genome (GRCh38.p14) using STAR (v2.7.11b, Dobin et al. 2013) with the Ensembl transcriptome annotation (v110) and the following non-default parameters `--alignEndsType EndToEnd`, `--outFilterMatchNminOverLread 0.9`, `--outSAMstrandField intronMotif`, `--outSAMattributes All`, `--outSAMtype BAM SortedByCoordinate`, `--seedSearchStartLmaxOverLread 0.5`, `--seedSearchStartLmax 20`, `--twopassMode Basic`. Genomic coordinates of 3' UTRs and coding sequences were downloaded from Ensembl Biomart (v110) for all protein-coding transcripts and filtered for transcript support level 1 or 2 or for the presence of their intron chain in the RefSeq annotation (`vGCF_000001405.40-RS_2023_10`).

### 4 Supplementary Tables

#### 4.1 Mapping statistics Ribo-seq data

| Sample | Total reads | Aligned | Aligned % | Uniq aligned | Uniq aligned % | Avg. mapped len | Annotated splices | Mismatch rate % |
|---|---|---|---|---|---|---|---|---|
| SRR10533453 | 2.407 | 2.009 | 83.480 | 1.046 | 43.480 | 29.510 | 0.162 | 0.460 |
| SRR10533454 | 44.084 | 38.132 | 86.500 | 21.729 | 49.290 | 29.620 | 4.054 | 0.780 |
| SRR10533455 | 40.297 | 34.144 | 84.730 | 17.073 | 42.370 | 29.610 | 2.725 | 0.490 |
| SRR10533456 | 38.349 | 35.468 | 92.490 | 21.411 | 55.830 | 29.750 | 3.394 | 0.270 |
| SRR10533457 | 37.477 | 34.770 | 92.770 | 21.615 | 57.670 | 29.440 | 3.414 | 0.270 |
| SRR10533458 | 41.920 | 36.853 | 87.920 | 22.292 | 53.180 | 29.350 | 3.567 | 0.300 |
| SRR10533459 | 35.910 | 33.663 | 93.740 | 13.750 | 38.290 | 29.000 | 2.163 | 0.370 |
| SRR10533460 | 35.633 | 32.009 | 89.830 | 11.399 | 31.990 | 29.590 | 1.761 | 0.370 |
| SRR10533461 | 21.238 | 18.893 | 88.960 | 8.600 | 40.490 | 29.330 | 1.296 | 0.390 |
| SRR10533462 | 29.909 | 26.093 | 87.240 | 8.211 | 27.450 | 28.580 | 1.341 | 0.900 |
| SRR10533463 | 55.638 | 49.527 | 89.020 | 15.315 | 27.530 | 30.120 | 2.459 | 0.410 |
| SRR10533464 | 30.649 | 24.209 | 78.990 | 9.019 | 29.430 | 28.910 | 1.322 | 0.500 |
| SRR10533465 | 41.252 | 38.511 | 93.350 | 13.837 | 33.540 | 28.650 | 1.994 | 0.400 |
| SRR10533466 | 39.912 | 35.797 | 89.690 | 16.173 | 40.520 | 29.110 | 2.462 | 0.380 |
| SRR10533467 | 32.341 | 29.639 | 91.640 | 10.351 | 32.000 | 28.780 | 1.444 | 0.270 |
| SRR10533468 | 24.664 | 23.244 | 94.240 | 6.190 | 25.100 | 28.450 | 0.867 | 0.330 |
| SRR10533469 | 44.206 | 38.939 | 88.080 | 18.246 | 41.270 | 28.750 | 2.736 | 0.300 |
| SRR11294608 | 34.789 | 27.581 | 79.290 | 7.934 | 22.810 | 31.590 | 1.464 | 1.970 |
| SRR11294609 | 34.477 | 27.312 | 79.210 | 8.103 | 23.500 | 31.480 | 1.608 | 2.060 |
| SRR11294610 | 29.093 | 21.678 | 74.510 | 5.563 | 19.120 | 31.670 | 0.753 | 1.960 |
| SRR11294611 | 42.023 | 31.655 | 75.320 | 8.452 | 20.110 | 31.480 | 1.546 | 2.290 |
| SRR8590754 | 17.254 | 14.260 | 82.650 | 7.212 | 41.800 | 28.980 | 0.814 | 1.610 |
| SRR8590755 | 17.110 | 14.469 | 84.560 | 7.176 | 41.940 | 28.980 | 0.842 | 1.620 |
| SRR8590756 | 15.321 | 13.704 | 89.440 | 5.831 | 38.060 | 29.230 | 0.630 | 1.620 |
| SRR8590757 | 15.191 | 13.545 | 89.170 | 5.772 | 38.000 | 29.210 | 0.629 | 1.620 |
| SRR8590758 | 18.026 | 15.782 | 87.550 | 5.839 | 32.390 | 29.280 | 0.629 | 2.100 |
| SRR8590759 | 18.038 | 15.743 | 87.280 | 5.852 | 32.440 | 29.300 | 0.616 | 2.090 |
| SRR8590766 | 28.658 | 25.183 | 87.880 | 8.224 | 28.700 | 29.650 | 0.819 | 1.600 |
| SRR8590767 | 29.121 | 25.732 | 88.360 | 8.329 | 28.600 | 29.650 | 0.805 | 1.590 |
| SRR8590768 | 36.289 | 32.328 | 89.090 | 12.656 | 34.880 | 29.050 | 1.463 | 1.710 |
| SRR8590769 | 36.896 | 33.562 | 90.960 | 12.797 | 34.680 | 29.070 | 1.449 | 1.680 |
| SRR8590778 | 32.271 | 28.487 | 88.270 | 11.160 | 34.580 | 29.460 | 1.107 | 1.350 |
| SRR8590779 | 32.206 | 27.495 | 85.370 | 11.062 | 34.350 | 29.470 | 1.113 | 1.340 |
| SRR8590780 | 7.716 | 6.675 | 86.520 | 2.930 | 37.970 | 29.210 | 0.271 | 2.450 |
| SRR8590781 | 7.241 | 6.539 | 90.320 | 2.792 | 38.560 | 29.380 | 0.251 | 1.660 |
| SRR8590782 | 7.624 | 6.884 | 90.290 | 2.931 | 38.450 | 29.180 | 0.271 | 2.470 |
| SRR8590783 | 7.294 | 6.308 | 86.500 | 2.851 | 39.100 | 29.350 | 0.255 | 1.630 |
| SRR8590790 | 47.881 | 40.691 | 84.980 | 19.535 | 40.800 | 29.490 | 2.002 | 1.820 |
| HCT_N_AID_UPF1_0h_IAA_1 | 9.039 | 8.833 | 97.710 | 4.557 | 50.410 | 28.030 | 0.626 | 0.620 |
| HCT_N_AID_UPF1_0h_IAA_2 | 19.819 | 18.531 | 93.510 | 9.805 | 49.480 | 27.650 | 1.379 | 0.680 |
| HCT_N_AID_UPF1_0h_IAA_3 | 18.632 | 18.190 | 97.620 | 8.888 | 47.700 | 27.940 | 1.280 | 0.620 |
| HCT_N_AID_UPF1_12h_IAA_1 | 30.060 | 29.711 | 98.840 | 18.644 | 62.020 | 28.110 | 2.648 | 0.520 |
| HCT_N_AID_UPF1_12h_IAA_2 | 35.302 | 34.623 | 98.070 | 21.528 | 60.980 | 28.000 | 3.104 | 0.560 |
| HCT_N_AID_UPF1_12h_IAA_3 | 7.373 | 7.100 | 96.300 | 3.024 | 41.010 | 27.740 | 0.413 | 0.720 |

**Table S1.** STAR alignment statistics for glioblastoma (PRJNA591767), breast cancer (PRJNA523167) and UPF1-degradation (E-MTAB-13837) Ribo-seq data.

| Sample | Total seqs | M Unique Reads | % Pass Dedup |
|---|---|---|---|
| HCT_N_AID_UPF1_0h_IAA_1 | 2.261 | 5.627 | 22.560 |
| HCT_N_AID_UPF1_0h_IAA_2 | 4.854 | 12.104 | 25.710 |
| HCT_N_AID_UPF1_0h_IAA_3 | 4.395 | 10.963 | 22.310 |
| HCT_N_AID_UPF1_12h_IAA_1 | 9.257 | 18.900 | 31.210 |
| HCT_N_AID_UPF1_12h_IAA_2 | 10.671 | 22.303 | 30.840 |
| HCT_N_AID_UPF1_12h_IAA_3 | 1.501 | 4.342 | 17.170 |

**Table S2.** Deduplication statistics for UPF1-degradation (E-MTAB-13837) Ribo-seq data.

### 5 Supplementary Figures

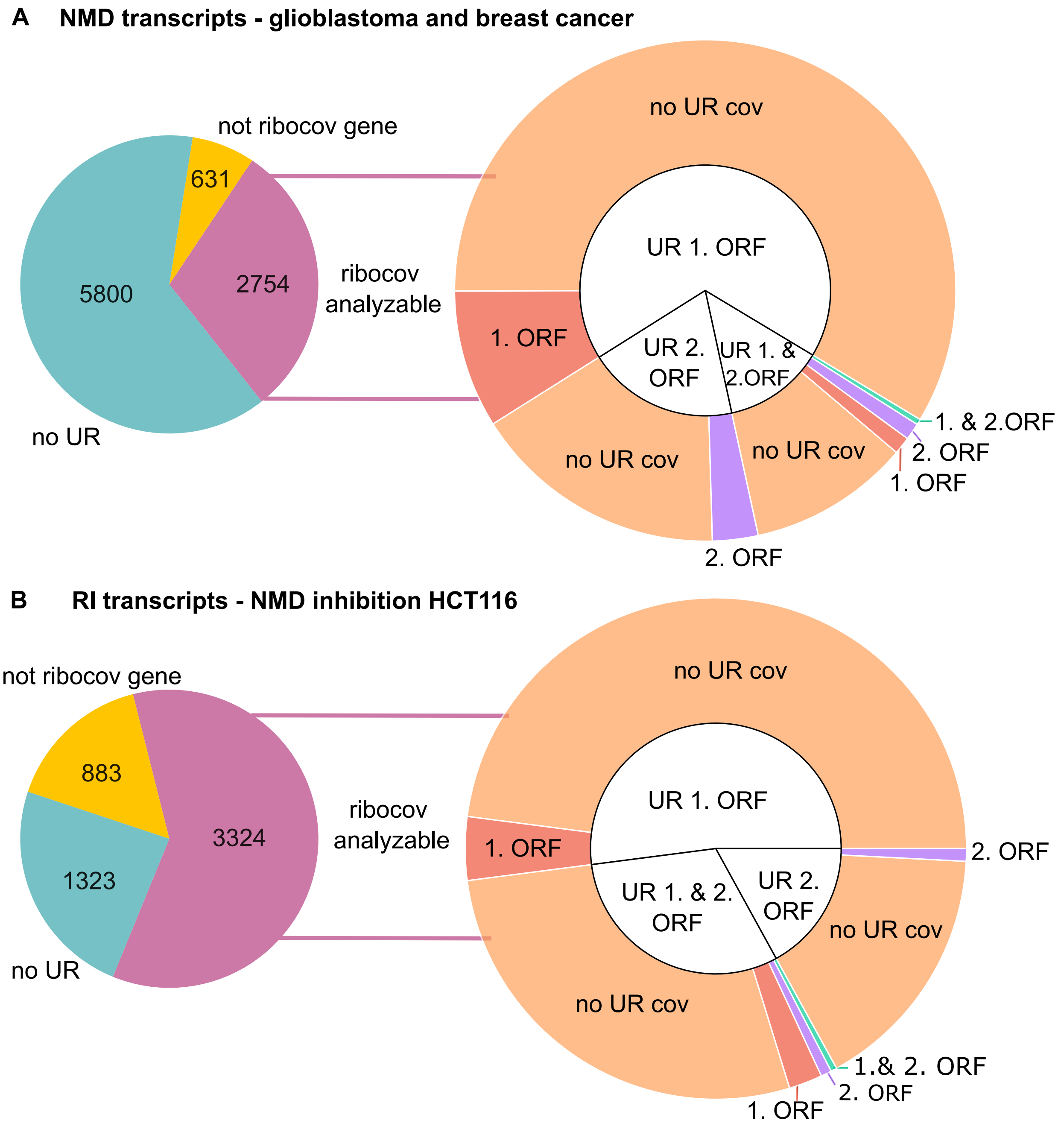


**Fig. 1. A**: Left: Pie chart indicating the number of ribo-cov analyzable NMD candidate Split-ORF transcripts with the glioblastoma and breast cancer samples. (Right) Sunburst chart of ribo-cov NMD candidate Split-ORF transcripts by unique region position for the glioblastoma and breast cancer samples. **B**: Left: Pie chart indicating the number of ribo-cov analyzable RI candidate Split-ORF transcripts with the HCT116 NMD inhibited samples. (Right) Sunburst chart of ribo-cov RI candidate Split-ORF transcripts by unique region position for the HCT116 NMD inhibited samples.

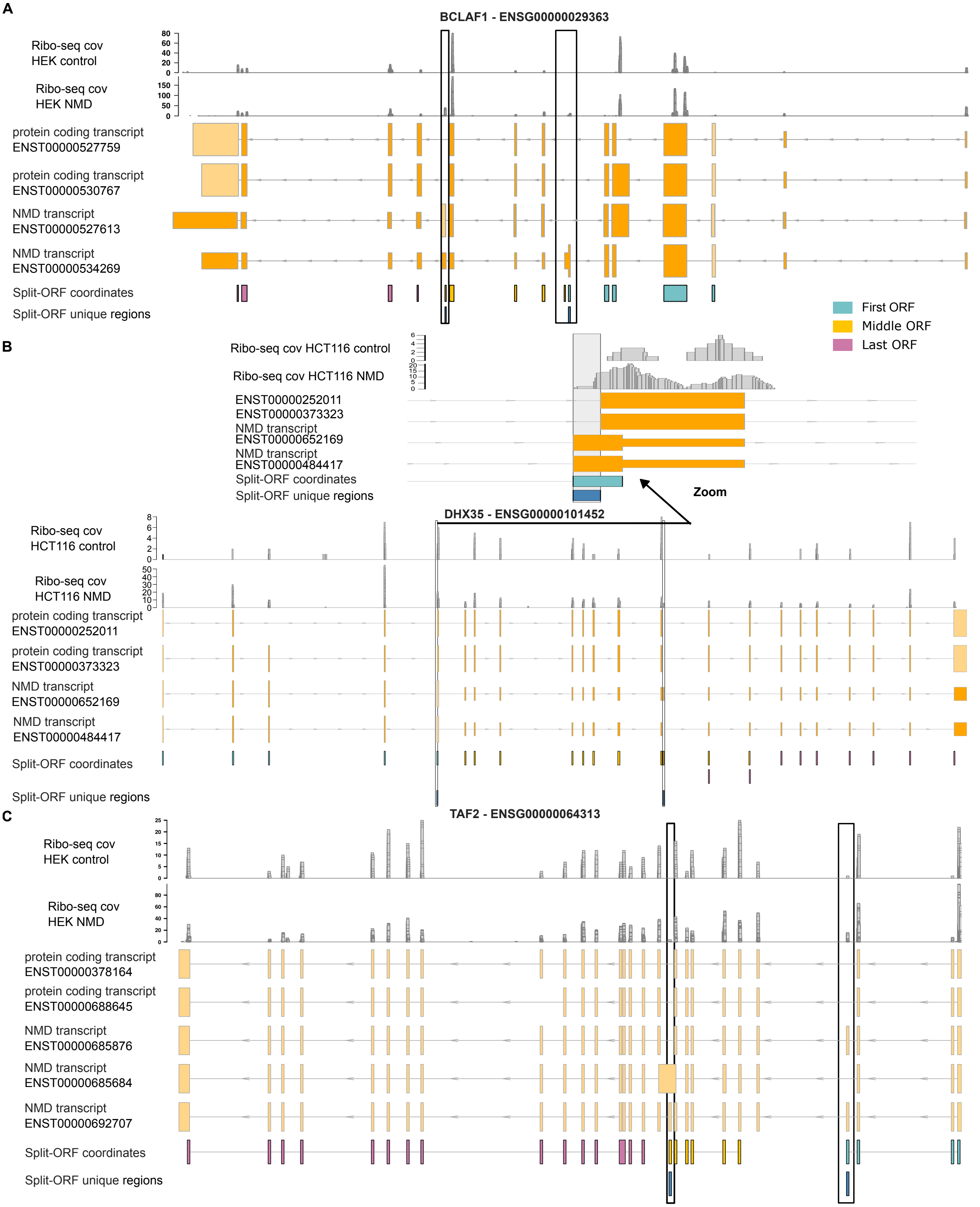


**Fig. 2. Ribo-seq coverage in both Split-ORF unique regions increases upon NMD inhibition in HCT116 cells.**. The y-axis of the Ribo-seq coverage is given in read depth per base. **A**: Example of Ribo-seq coverage in *BCLAF1* transcripts and unique regions of NMD candidate Split-ORF transcripts in control and NMD inhibition. **B**: Example of Ribo-seq coverage in *DHX35* transcripts and unique regions of NMD candidate Split-ORF transcripts in control and NMD inhibition. **C**: Example of Ribo-seq coverage in *TAF2* transcripts and unique regions of NMD candidate Split-ORF transcripts in control and NMD inhibition.

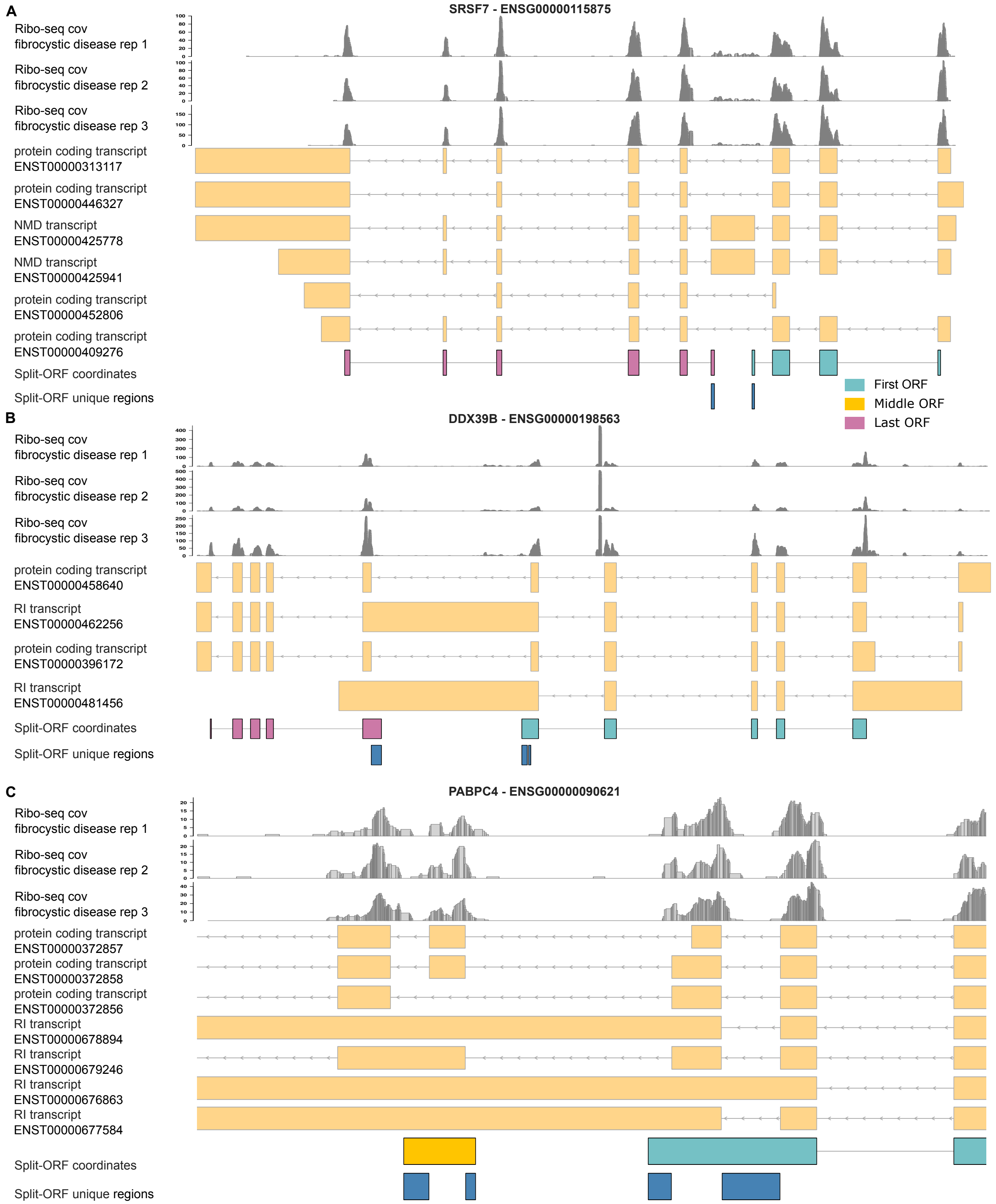


**Fig. 3. Ribo-seq coverage in both Split-ORF unique regions in fibrocystic disease cells.** The y-axis of the Ribo-seq coverage is given in read depth per base. **A**: Example of Ribo-seq coverage in *SRSF7* transcripts and unique regions of NMD candidate Split-ORF transcripts. **B**: Example of Ribo-seq coverage in *DDX39B* transcripts and unique regions of RI candidate Split-ORF transcript. **C**: Example of Ribo-seq coverage in *PABPC4* transcripts and unique regions of RI candidate Split-ORF transcripts.
